# Prevalence of Avian Influenza Virus in Synanthropic Birds Associated with an Outbreak of Highly Pathogenic Strain EA/AM H5N1

**DOI:** 10.1101/2023.11.08.565892

**Authors:** Jourdan M. Ringenberg, Kelsey Weir, Lee Humberg, Carl Voglewede, Mitch Oswald, J. Jeffrey Root, Krista Dilione, Evan Casey, Michael Milleson, Timothy Linder, Julianna Lenoch

## Abstract

The 2022 – 2023 highly pathogenic avian influenza (HPAI) virus outbreak of H5N1 Eurasian lineage goose/Guangdong clade 2.3.4.4b is the largest in North American history and has significantly impacted wild bird populations and domestic poultry across the United States. Synanthropic birds may play an important role in transmitting the virus laterally to other wild bird species and domestic poultry. Understanding the prevalence of HPAI H5N1 in different avian orders may help inform management decisions and potential risk factors for both wild and domestic bird populations. Following the confirmation of infection of HPAI H5N1 in domestic poultry at two commercial premises in IN, USA, we sampled and tested 266 synanthropic avian species within the Columbiformes and Passeriformes orders and found no detection of the virus at either location. Additionally, laboratories within the National Animal Health Laboratory Network were queried for influenza Type A rRT-PCR assay test results from morbidity and mortality events in wild birds, consisting of 10,786 birds tested across eight orders and 1,666 avian influenza virus detections. Query results were assessed by taxonomic groups for viral prevalence and suggested that the virus most often was observed in predatory and scavenging birds. Although detections were found in non-predatory synanthropic birds including the orders Columbiformes, Galliformes, and Passeriformes, the risk of transmission from and between these groups appears comparatively low, with apparent prevalence rates of 0.0090, 0.0688, and 0.0147, respectively. The highest prevalence was observed in raptors (0.2514), with prevalence rates in exclusively scavenging *Cathartidae* reaching up to 0.5333. There is strong evidence that consumption of infected tissues is a key pathway for transmission of avian influenza viruses. Understanding the impact of the 2022 – 2023 HPAI outbreak in wild bird populations can provide pertinent information on viral transmission, disease ecology, and risk to humans and agriculture.

## Introduction

The outbreak of highly pathogenic avian influenza (HPAI) H5N1 Eurasian lineage goose/Guangdong (Gs/GD) clade 2.3.4.4b virus (hereafter H5N1) throughout 2022 and 2023 is the largest in North American history and has impacted wild bird populations and domestic poultry significantly across the continent. The first known infection of H5N1 in North America occurred in a wild great black-backed gull (*Larus marinus*) from Newfoundland and Labrador Province, Canada, in November 2021 [1]. In January 2022, H5N1 was reported in apparently healthy wild waterfowl from NC and SC, USA, and since has been detected in wild birds in 49 United States (U.S.) states [2]. As HPAI H5Nx subtypes continue to circulate throughout Eurasia and the Americas [3,4], the migratory nature of wild birds introduces the risk of recombination and reassortment and the introduction of new strains into North America [5,6,7]. Understanding the prevalence in wild bird species can help inform management decisions for wild bird populations and the commercial poultry industry.

The avian orders Anseriformes (ducks, geese, and swans) and Charadriiformes (shorebirds, gulls, and terns) act as the primary reservoir hosts of avian influenza (AI) viruses in the wild [8,9]. While waterfowl play a significant role in the transmission of AI viruses due to their gregarious, migratory nature and their potential for significant viral shedding, evidence has shown that they often present as asymptomatic and survive viral infection [10,11]. Significant research has been conducted on AI viruses in Anseriformes and Charadriiformes, which predominantly replicate AI viruses in the intestinal and respiratory tract and can readily transmit AI viruses by the oral – fecal route to other avifauna that share water resources [6,12]. While methods of viral transmission are well understood in these orders, less is known about the role alternative hosts play in transmitting AI viruses across the landscape during an HPAI virus outbreak. Understanding the viral prevalence of different orders may help identify areas with greater risk of HPAI virus infection to alternative avian hosts, threatened and endangered species, and domestic poultry.

Although the previous North American outbreak of HPAI Eurasian H5 viruses in 2014 – 2015 caused mortality in some wild bird species, the impact was less severe than the 2022 – 2023 outbreak. Between December 2014 and June 2015, 98 birds tested positive for HPAI viruses out of approximately 7,084 wild birds sampled: 75 from apparently healthy waterfowl, 16 from mortality events involving snow geese (*Chen caerulescens*) and ringed-necked ducks (*Aythya collaris*), and seven captive raptor mortalities [13]. Conversely, with over 7,400 confirmed HPAI H5Nx detections in the USA since January 2022 in over 150 wild bird species across numerous avian orders, the impact of this outbreak on wild bird populations is much greater [2].

Domestic poultry populations in the USA also have been substantially impacted following the initial detection of HPAI H5N1 in a commercial turkey facility in IN in February 2022. Detections in domestic poultry (commercial and backyard flocks) have occurred alongside wild bird detections throughout the course of the 2022 – 2023 outbreak with confirmed infections in 47 states [14]. Initial genetic sequencing conducted by the U.S. Department of Agriculture (USDA) National Veterinary Services Laboratories (NVSL) suggests that most poultry detections have wild bird origins with a minority occurring by lateral transmission [15]. Thus, the concern from the commercial poultry industry is high and has triggered increased surveillance in wild birds since the beginning of the outbreak in 2022. Commercial poultry assessments investigating routes of transmission following previous HPAI outbreaks have identified high risk factors such as poor biosecurity practices and the movement of people, equipment, and domestic birds [16]. Exact mechanisms of H5N1 transmission from wild birds to poultry throughout 2022 – 2023 are speculative, but bridge hosts, which are non-maintenance host species that can transmit pathogens from reservoir species to domestic poultry through shared resources (e.g., water, crops, feed sources), could play a vital role [17,18]. Synanthropes, or species that are ecologically associated with human populations and regularly utilize anthropogenically modified environments, may act as bridge hosts [19]. Synanthropic species, such as those from the families *Columbidae* and *Passeriformes*, are often found in and around poultry facilities and thus are suspected as potential routes of HPAI virus transmission, as they could act to transport viruses between infected commercial premises. Expanding the lens to other synanthropic species and their potential role in transmission, particularly considering the 2022 – 2023 outbreak, can help better focus management resources and mitigate viral spread.

Known broadly for their synanthropic behavior, species in the order Columbiformes (doves and pigeons) often have been the subject of AI virus research [19] and investigated as potential bridge hosts in transmitting AI viruses between migratory birds and poultry or between poultry facilities during disease outbreaks [20]. Experimental infections of rock doves (*Columba livia;* often referred to as pigeons) have shown their role in AI virus transmission is likely via fomite or mechanical routes, and when they do shed virus, the quantities and time frames of shedding are limited [19,21]. While the risk for transmission to domestic poultry is low, there is evidence that some AI virus strains can spread from Columbiformes to other avian species and cause infection [19].

The order Passeriformes contains several families of birds that demonstrate synanthropic behavior, including *Corvidae* (crows, jays, magpies, and ravens), *Fringillidae* (finches), *Hirundinidae* (swallows), *Icteridae* (blackbirds and grackles), *Passeridae* (Old World sparrows), *Sturnidae* (starlings), and *Turdidae* (robins and thrushes) [19]. Many species within these families commonly are found on farms and have the potential to act as bridge hosts. Susceptibility to AI viruses has been shown both experimentally and in the wild in several Passeriformes species. In their review evaluating AI virus infection rates in wild birds globally, Caron, Cappelle, and Gaidet [22] calculated a 0.0206 prevalence rate for all Passeriformes tested; however, evidence of AI virus susceptibility differs between species.

Many *Corvidae* species are omnivorous, opportunistic foragers, and keen scavengers that commonly are attracted to carcasses accessible on farms. Studies evaluating both natural and experimental infections of HPAI viruses in *Corvidae* suggest they may play an important ecological and epidemiological role in HPAI H5 viruses. In South Korea in 2003 – 2004, H5N1 was detected in Korean magpies (*Pica pica sericea*) found dead at a poultry facility [23], and investigations of large-billed crow (*Corvus macrorhynchos)* mortalities closely associated with an H5N1 domestic poultry outbreak in Japan in 2004 demonstrated their susceptibility to infection [24]. Experimental inoculation of house crows (*Corvus splendens*) with H5N1 crow and chicken virus isolates caused clinical signs and mortalities in 66.7% and 50% of study animals, respectively [25], suggesting the potential for virus transmission between crows and poultry. Rooks (*Corvus frugilegus)* experimentally inoculated with HPAI H5 all seroconverted and shed virus with a 25% mortality rate [26]. Furthermore, an assessment of risk factors predicting H5N1 infections on poultry farms in Bangladesh identified house crows as the greatest risk factor for virus dispersal [27]. While susceptibility to H5N1 has been demonstrated in several cases, more research is warranted to determine the role *Corvidae* play in virus transmission.

Non-*Corvidae* species in the Passeriformes order often are colloquially referred to as songbirds, but distinct differences between them have important implications for HPAI virus susceptibility and transmission. Of species in the *Fringillidae* family, house finches (*Haemorhous mexicanus*) commonly display synanthropic behavior, yet the few assessments of their susceptibility to AI viruses have found low prevalence rates suggesting the risk of transmission is low [19]. While the insectivorous diet of *Hirundae* species could decrease their likelihood of interacting with poultry or shared resources [19], their global abundance and occupancy on farms stresses the importance of understanding their role in AI virus ecology [19]. Studies have demonstrated swallows’ susceptibility to AI viruses [28,29] and potential to act as bridge hosts [30,31]. Several *Icteridae* species are a common presence on farms, including the common grackle (*Quiscalus quiscula*), red-winged blackbird (*Agelaius phoeniceus*), and brown-headed cowbird (*Molothrus ater*) [19]. Results of AI virus transmission in *Icteridae* species are mixed, and more research is needed to better understand the role they play in spillover to poultry. Within the *Passeridae* family, sparrows are susceptible to many AI viruses of which they can shed high levels and transmit to poultry [19]. Two studies that experimentally inoculated (1) tree sparrows (*Spizelloides arborea*) with four HPAI H5Nx virus strains [32] and (2) house sparrows (*Passer domesticus*) with HPAI H5N1 [21] found both species to be highly susceptible. European starlings (*Sturnus vulgaris*), the most common, widespread synanthrope in the *Sturnidae* family, often flock to farms for food resources and nesting sites in groups so large that even small amounts of viral shedding by individuals collectively could cause AI virus spillover to poultry [19,33,34]. Starlings sampled and tested for AI viruses across 14 studies showed a 0.018 prevalence rate, but their role in transmission may be strain-dependent [19]. Within the *Turdidae* family, AI viruses were detected in American robin (*Turdus migratorius*) and Swainson’s thrush (*Catharus ustulatus*) at rates of 0.0376 and 0.0377, respectively, during a surveillance study conducted in passerines across the USA [35]. While an experimental study inoculated American robins with HPAI H5Nx viruses and found 0.8800 prevalence [36]. Ultimately, songbird susceptibility to AI viruses is variable, and more work is needed to evaluate the spillover risk to poultry.

The order Galliformes (pheasants, turkeys, peafowl, and quail) often exhibit synanthropic behavior and evidence has shown that many species in this family are susceptible to and can shed AI viruses [19]. Many Galliformes that have been studied are domesticated and raised in backyard or gamebird farms, and less is understood about the contact frequency between wild and domestic individuals and AI virus dynamics in wild Galliformes. Galliformes have the potential to act as bridge hosts, as agricultural areas may attract wild individuals searching for food resources or conspecifics [19]. A serosurvey in Italy of 219 free-living pheasants (*Phasianus colchicus*) found a 0.1230 prevalence rate but detected no antibodies to low-pathogenic avian influenza (LPAI) virus H5 subtypes [37]. A similar study of hunter-harvested, wild-captured bobwhite quail (*Colinus virginianus*) in TX, USA, found 1.4% positive and 7.6% suspect for AI viruses [38].

Feeding methods of avian scavengers and predators provide the opportunity for contact with HPAI virus-infected carcasses or prey. Susceptibility to HPAI viruses is high, and exposures and infections have been detected in Accipitriformes (hawks and eagles), Cathartiformes (New World vultures), Falconiformes (falcons), and Strigiformes (owls) [19]. Bertran et al. [39] confirmed both HPAI and LPAI virus transmission to Gyr-Saker hybrid falcons (*Falco rusticolus* x *Falco cherrua*) through the experimental ingestion of infected chickens. While conducting passive surveillance following the HPAI H5Nx outbreak in the USA in 2014 – 2015, Ip et al. [10] found raptors (hawks, eagles, and owls) to be particularly susceptible to HPAI H5 viruses, with an overall positivity rate of 52.4%. Hall et al. [40] found American kestrels (*Falco sparverius*) to be highly susceptible to H5N1 with 100% mortality rate of experimentally inoculated birds. However, other studies have noted low prevalence in raptor species. Findings in an examination of raptors in OK, USA, found only 0.0160 prevalence in red-tailed hawks (*Buteo jamaicensis*) [41]. Raptors that specifically scavenge or prey upon aquatic birds were screened for influenza A antibodies at wildlife rehabilitation centers in MN and VA, USA [42]. They found evidence of AI virus exposure in bald eagles (*Haliaeetus leucocephalus*; 5.1%), negligible evidence of exposure in peregrine falcons (*Falco peregrinus*; 0.2%), great horned owls (*Bubo virginianus*; 1.2%), and Cooper’s hawks (*Accipiter cooperii*; 1.0%), and zero evidence of exposure in vultures, concluding that bald eagles likely would be affected by HPAI viruses should one be detected in waterfowl. Regardless, there is strong historical evidence of susceptibility to highly pathogenic and other AI viruses in these orders and understanding their prevalence throughout the 2022 – 2023 outbreak can help add to the body of knowledge and provide management insight [43,44].

Our investigation focuses on synanthropic species submitted for HPAI testing as part of morbidity/mortality (M/M) investigations and commercial poultry facility sampling events during the 2022 – 2023 H5N1 outbreak. For the purposes of our study, synanthropic refers to terrestrial wild bird species that are (1) non-reservoir hosts of AI viruses, (2) associated ecologically with human populations, and (3) regularly utilize anthropogenically modified environments [19]. Waterfowl and other aquatic, coastal, and pelagic orders such as Pelecaniformes, Anseriformes, and Charadriiformes are excluded from this evaluation.

The objectives of this study were (1) to assess the presence of HPAI viruses in synanthropic birds captured around H5N1-positive commercial poultry premises in response to the initial detection in domestic poultry in the USA and (2) to evaluate the prevalence of AI viruses in synanthropic bird orders during an HPAI outbreak. To address the first objective, we initiated a surveillance project to sample synanthropic bird species around HPAI-affected commercial poultry premises and tested for the presence of HPAI. To address the second objective, we evaluated data from the National Animal Health Laboratory Network (NAHLN) on wild bird species submitted for AI virus diagnostic testing as part of morbidity/mortality (M/M) investigations. In this study, we report results from the targeted surveillance project, compare prevalence rates of AI viruses in several avian orders submitted from M/M investigations from February 2022 to March 2023, and provide the total number of HPAI H5Nx positive birds confirmed by the NVSL from avian orders of interest.

## Materials and Methods

### Targeted Surveillance

Synanthropic bird species were sampled at two adjacent commercial domestic turkey farms with confirmed HPAI H5N1 in Dubois County, IN, USA, in February 2022. Samples were collected in accordance with the USDA Wild Bird Avian Influenza Surveillance Field Procedures Manual (Summer FY2022 to Winter FY2023) and within the guidelines and regulations set forth by the U.S. Fish and Wildlife Service (USFWS) under permit number MB124992. All samples were collected with the permission of the farm owners. Sampling of wild birds began approximately two weeks following virus detection and the initiation of poultry depopulation. A clean and dirty line was established on both premises, requiring all people, vehicles, supplies, and equipment to be fully cleaned and disinfected prior to crossing from the dirty side to the clean side. Traps were deployed to target European starlings, house sparrows, and rock doves. Five trap designs were used: custom three-hole wooden nest box traps composed of vertically stacked Sherman traps (H. B. Sherman Traps, Inc., Tallahassee, FL, USA); custom made PVC single hole nest box traps with PVC caps and a single catch trap door (Van Ert Enterprises, Decatur, IA, USA); custom portable single-axle trailer drop-in starling decoy traps; baited walk-in traps with funnels; and decoy, walk-in pigeon traps (Tomahawk Live Trap, Hazelhurst, WI, USA). Traps were set within the perimeter of the infected farms on the clean side of the line and placed around poultry barns, grain bins, feed silos, other farm structures, and suspected avian movement corridors on the edges of natural or agriculturally modified habitat. Traps were set every morning on each site and checked within 24 hours for a total of 18 days. Traps were baited with commercial bird seed, dry cat food, and corn. Traps were disinfected with Virkon™ S (LANXESS, Pittsburgh, PA, USA) before transferring to a new location.

All captured species were identified by field biologists. Upon capture, birds were immediately euthanized via cervical dislocation and subsequently sampled. Oropharyngeal and cloacal swabs (Harmony Lab and Safety Supplies, Grove Garden, CA, USA) were collected from all captured birds. Both swabs were pooled into a single tube containing 1.5 mL of PrimeStore® Molecular Transport Medium (MTM; EKF Diagnostic, Barleben, Germany) and were shipped to the Veterinary Diagnostic Laboratory at Colorado State University within three days to maintain sample integrity. Nucleic acids were extracted from the samples following standard extraction protocols, and a general influenza Type A rRT-PCR assay targeting the conserved region of the avian influenza matrix gene was performed [45,46]. Prevalence rates were calculated for each species sampled and tested.

### Morbidity and Mortality Investigations

Morbidity and mortality (M/M) investigations were conducted across numerous species of birds that appeared sick, moribund, or dead due to suspected exposure to HPAI H5N1 within the conterminous U.S. and Alaska throughout the 2022 – 2023 outbreak. Tracheal and cloacal swabs, whole carcasses, or tissue samples were collected opportunistically by state agencies, federal agencies, or rehabilitation facilities. Sampling methodologies may have differed depending on the collecting state, agency, or facility in terms of the number of birds sampled at each M/M event and type(s) of samples collected. Samples were submitted to labs in the NAHLN for diagnostic testing, which included a general influenza Type A rRT-PCR assay for all samples, and any samples with a resulting non-negative cycle threshold (Ct) value were further tested using an H5 rRT-PCR subtyping assay [45,46].

We queried all laboratories in the NAHLN and provided a standardized spreadsheet to be completed with a list of species across multiple taxonomic groups. We focused on groups that most commonly exhibit synanthropic behavior but excluded known reservoir hosts and other waterfowl species. Labs recorded the number of each species tested from 1 February 2022 to 31 March 2023, and the resultant number of non-negative samples as determined by the general influenza Type A rRT-PCR assay. Responses were compiled to calculate the prevalence of AI viruses in each sampled species, and species were grouped by order and family. Known captive and domestic birds were excluded from the dataset.

### Confirmatory Testing of HPAI H5 Detections

Lastly, we report the total number of H5Nx positive samples for synanthropic orders of interest from the wild bird HPAI detection dataset [2]. These samples had previously undergone confirmatory testing at the NVSL, which included an rRT-PCR assay targeting Eurasian lineage Gs/GD H5 clade 2.3.4.4b (SEPRL; Real-Time RT-PCR Assay for the Detection of Goose/Guangdong lineage Influenza A subtype H5, clade 2.3.4.4; NVSL-WI-1732), as well as an N1 subtyping rRT-PCR assay (SEPRL; Real-Time RT-PCR Assay for the Detection of Eurasian-lineage Influenza A Subtype N1; NVSL-WI-1768). Samples submitted to the NVSL for confirmatory testing included those submitted as part of M/M investigations as well as samples collected from apparently healthy birds as part of targeted surveillance programs. Samples submitted from birds belonging to the orders Anseriformes, Charadriiformes, Pelecaniformes, Suliformes, and Gruiformes were removed from the dataset.

## Results

### Targeted Surveillance

Samples were obtained from a total of 266 wild synanthropic birds across eight species from two adjacent commercial turkey farms with confirmed HPAI H5N1 in Dubois County, IN (Table 1). None of the 266 individuals tested positive for influenza A virus by rRT-PCR from pooled cloacal and oral swabs, resulting in zero prevalence of AI virus in the sample. Samples were obtained from the families *Columbidae* (44), *Icteridae* (81), *Passeridae* (89), and *Sturnidae* (52).

**Table 1.** Number of synanthropic bird species sampled and prevalence of avian influenza virus at HPAI – affected commercial farms in Dubois Co, IN.

| Family | Species | Number of Birds Sampled | Influenza Type A rRT-PCR Detections (N Positive) | Prevalence |
| --- | --- | --- | --- | --- |
| *Columbidae* | Eurasian collared-dove | 1 | 0 | 0 |
|  | Mourning dove | 3 | 0 | 0 |
|  | Rock dove | 40 | 0 | 0 |
|  |  | 44 | 0 | 0 |
| *Icteridae* | Brown-headed cowbird | 51 | 0 | 0 |
|  | Common grackle | 17 | 0 | 0 |
|  | Red-winged blackbird | 13 | 0 | 0 |
|  |  | 81 | 0 | 0 |
| *Passeridae* | House sparrow | 89 | 0 | 0 |
|  |  | 89 | 0 | 0 |
| *Sturnidae* | European starling | 52 | 0 | 0 |
|  |  | 52 | 0 | 0 |
| Total |  | 266 | 0 | 0 |

### Morbidity and Mortality Investigations

Out of the 48 labs queried in the NAHLN, 32 labs (67%) provided AI virus diagnostic testing data broken down by individual species. Of these labs, a total of 10,786 birds were tested and 1,666 AI virus detections were observed (prevalence of 0.1545; see Table S1 in the Supplementary Material for a comprehensive list of all species tested). Prevalence rates were highest in Cathartiformes followed by Strigiformes, Accipitriformes, Falconiformes, Galliformes, Passeriformes, and Columbiformes (Figure 1).

**Figure 1.**
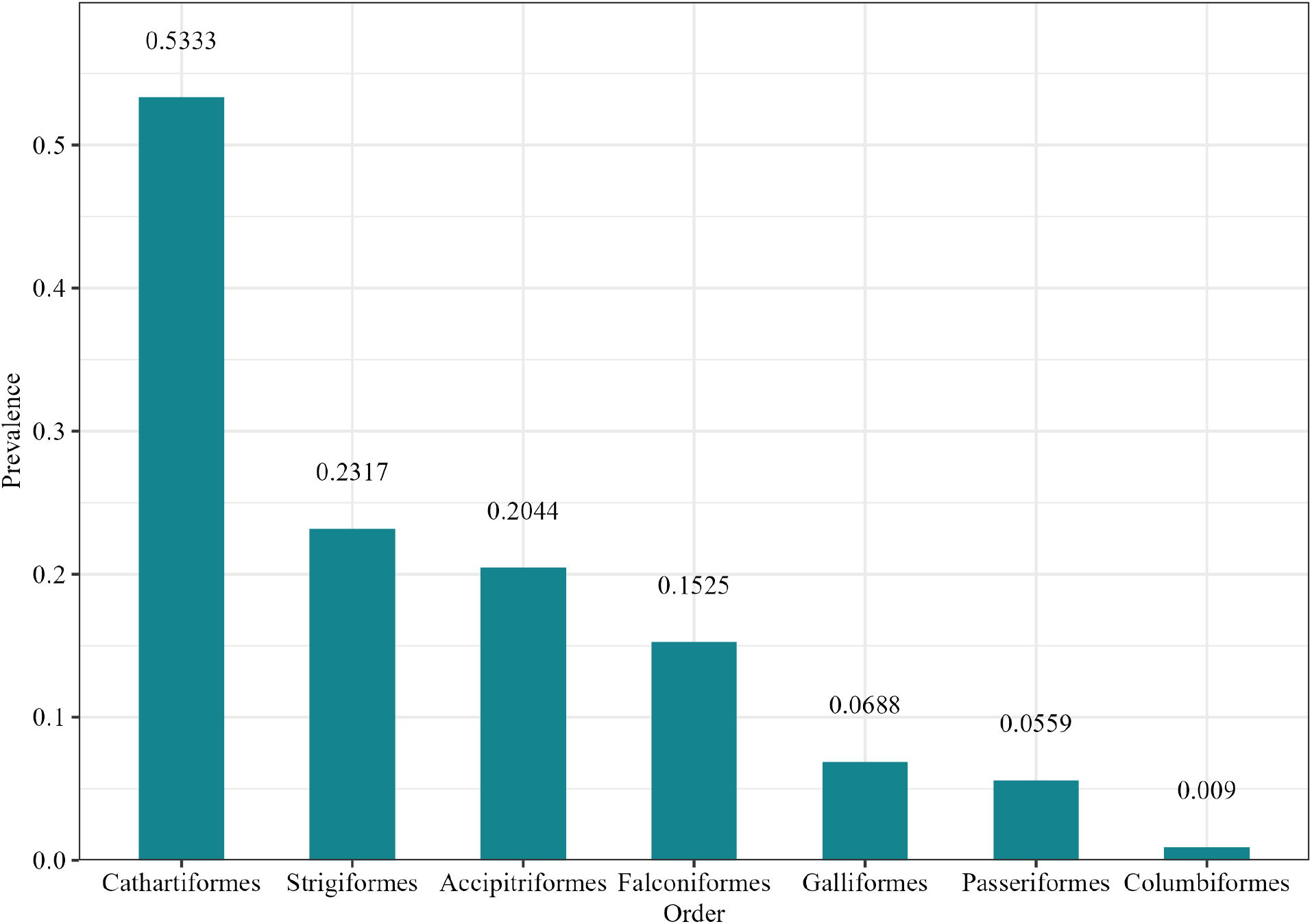
Prevalence of avian influenza A virus. Detections of AI viruses in avian orders submitted to the NAHLN as part of morbidity/mortality investigations from 1 February 2022 to 31 March 2023.

### Pigeons, Doves: Order Columbiformes, Family *Columbidae*

Out of the 443 samples collected from the family *Columbidae*, four tested positive for AI viruses, resulting in a prevalence of 0.0090 (Table 2). Mourning doves (*Zenaida macroura*) and rock doves accounted for 76% of *Columbidae* samples and all AI virus detections, with a slightly higher prevalence rate in mourning doves (0.0217) than rock doves (0.0082).

**Table 2.**
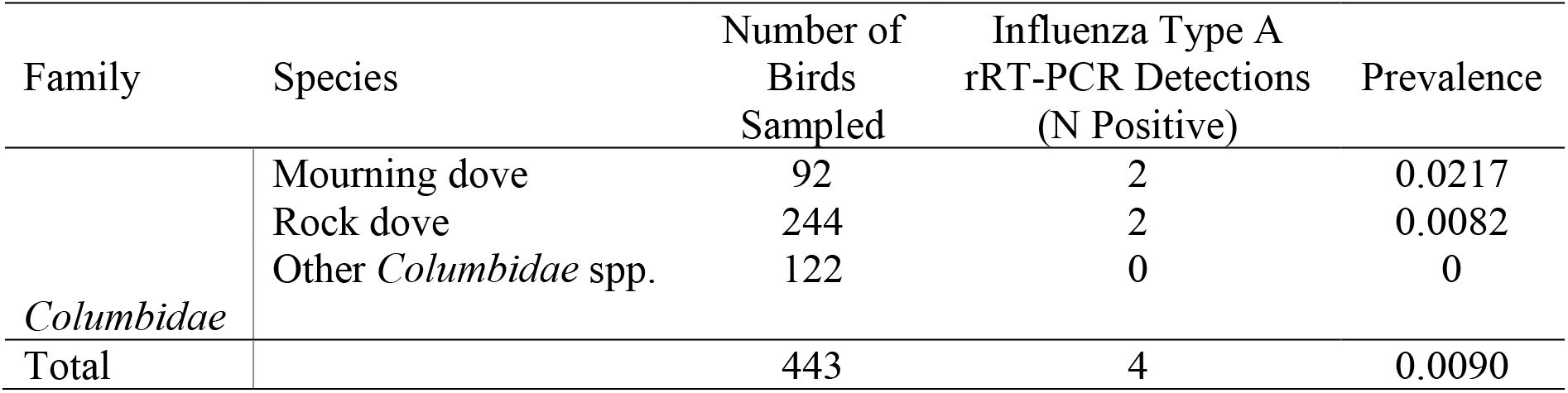
Avian influenza A virus detections in *Columbidae* morbidity/mortality submissions as reported by diagnostic laboratories in the NAHLN from 1 February 2022 to 31 March 2023.

### Songbirds: Orders Passeriformes and Piciformes

A total of 889 samples were obtained from the orders Passeriformes and Piciformes, 13 of which tested positive for AI viruses, resulting in a total prevalence of 0.0150 (Table 3). Of the families tested, AI virus was detected in *Fringillidae*, *Hirundinidae*, *Icteridae*, *Passerellidae*, *Passeridae*, and *Turdidae*. Prevalence was highest in *Hirundinidae* (0.1429), with five total detections in swallow species (*Tachycineta bicolor* and *Tachycineta thalassina). Fringillidae* had a prevalence of 0.0426, with one detection each in an American goldfinch (*Spinus tristis*) and pine grosbeak (*Pinicola enucleator*). While prevalence was highest in the pine grosbeak (0.2500), the sample size was small with only four birds tested. *Passerellidae* had a prevalence of 0.0333, with one detection in a dark-eyed junco (*Junco hyemalis*). *Icteridae* yielded a prevalence of 0.0250, with one detection each in a boat-tailed grackle (*Quiscalus major*), a common grackle, and a red-winged blackbird. Lowest prevalence rates were observed in the families *Passeridae* (0.0061), with one house sparrow detection, and *Turdidae* (0.0040), with one American robin detection.

**Table 3.**
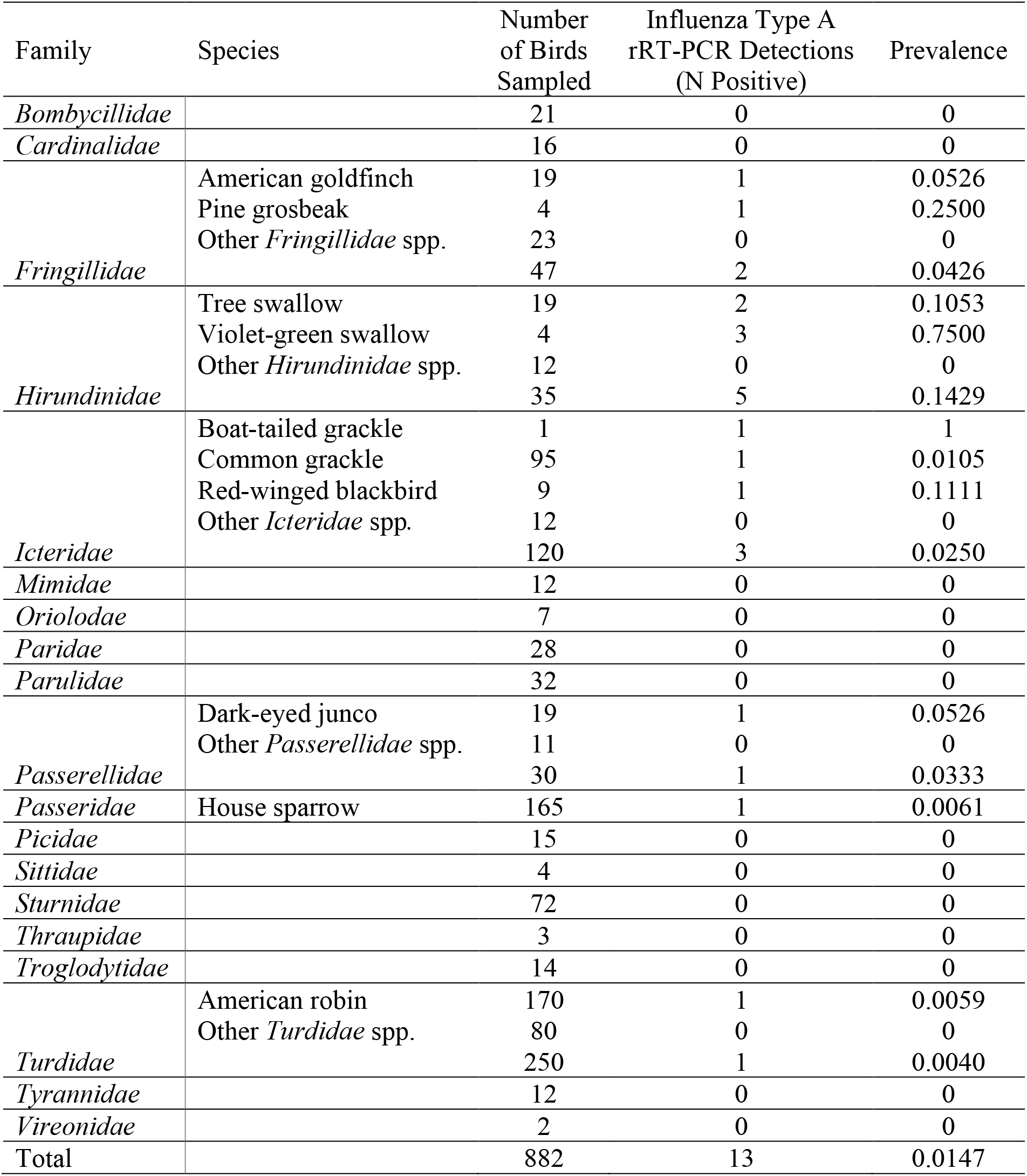
Avian influenza A virus detections in songbird morbidity/mortality submissions as reported by diagnostic laboratories in the NAHLN from 1 February 2022 to 31 March 2023.

| Family | Species | Number of Birds Sampled | Influenza Type A rRT-PCR Detections (N Positive) | Prevalence |
| --- | --- | --- | --- | --- |
| *Bombycillidae* |  | 21 | 0 | 0 |
| *Cardinalidae* |  | 16 | 0 | 0 |
| *Fringillidae* | American goldfinch | 19 | 1 | 0.0526 |
|  | Pine grosbeak | 4 | 1 | 0.2500 |
|  | Other *Fringillidae* spp. | 23 | 0 | 0 |
|  |  | 47 | 2 | 0.0426 |
| *Hirundinidae* | Tree swallow | 19 | 2 | 0.1053 |
|  | Violet-green swallow | 4 | 3 | 0.7500 |
|  | Other *Hirundinidae* spp. | 12 | 0 | 0 |
|  |  | 35 | 5 | 0.1429 |
| *Icteridae* | Boat-tailed grackle | 1 | 1 | 1 |
|  | Common grackle | 95 | 1 | 0.0105 |
|  | Red-winged blackbird | 9 | 1 | 0.1111 |
|  | Other *Icteridae* spp. | 12 | 0 | 0 |
|  |  | 120 | 3 | 0.0250 |
| *Mimidae* |  | 12 | 0 | 0 |
| *Oriolidae* |  | 7 | 0 | 0 |
| *Paridae* |  | 28 | 0 | 0 |
| *Parulidae* |  | 32 | 0 | 0 |
| *Passerellidae* | Dark-eyed junco | 19 | 1 | 0.0526 |
|  | Other *Passerellidae* spp. | 11 | 0 | 0 |
|  |  | 30 | 1 | 0.0333 |
| *Passeridae* | House sparrow | 165 | 1 | 0.0061 |
| *Picidae* |  | 15 | 0 | 0 |
| *Sittidae* |  | 4 | 0 | 0 |
| *Sturnidae* |  | 72 | 0 | 0 |
| *Thraupidae* |  | 3 | 0 | 0 |
| *Troglodytidae* |  | 14 | 0 | 0 |
| *Turdidae* | American robin | 170 | 1 | 0.0059 |
|  | Other *Turdidae* spp. | 80 | 0 | 0 |
|  |  | 250 | 1 | 0.0040 |
| *Tyrannidae* |  | 12 | 0 | 0 |
| *Vireonidae* |  | 2 | 0 | 0 |
| Total |  | 882 | 13 | 0.0147 |

### Crows, Ravens, Jays, and Magpies: Order Passeriformes, Family *Corvidae*

Of the 531 *Corvidae* tested, 66 were positive for AI viruses, resulting in a total prevalence of 0.1240 (Table 4). Prevalence was highest in common ravens (*Corvus corax*; 0.2358), followed by fish crows (*Corvus ossifragus*; 0.2083), magpies (*Pica* spp; 0.1429), and American crows (*Corvus brachyrhynchos*; 0.0997).

**Table 4.** Avian influenza A virus detections in *Corvidae* morbidity/mortality submissions as reported by diagnostic laboratories in the NAHLN from 1 February 2022 to 31 March 2023.

| Family | Species | Number of Birds Sampled | Influenza Type A rRT-PCR Detections (N Positive) | Prevalence |
| --- | --- | --- | --- | --- |
|  | American crow | 301 | 30 | 0.0997 |
|  | Common raven | 106 | 25 | 0.2358 |
|  | *Pica* spp. | 42 | 6 | 0.1429 |
|  | Fish crow | 24 | 5 | 0.2083 |
|  | Other *Corvidae* spp. | 59 | 0 | 0 |
| *Corvidae* |  |  |  |  |
| Total |  | 532 | 66 | 0.1240 |

### Raptors: Orders Accipitriformes, Cathartiformes, Strigiformes, and Falconiformes

Of the 5,306 raptor samples submitted for testing, 1,334 were positive for AI viruses, resulting in a total prevalence of 0.2514 (Table 6). Prevalence was highest in the *Cathartidae* family (0.5333), followed by *Strigidae* (0.2318), *Accipitridae* (0.2044), *Falconidae* (0.1525), and *Pandionidae* (0.0488). With a sample size of one, the prairie falcon (*Falco mexicanus*) had the highest prevalence (1.000) of all raptor species. The next highest prevalence rates were from black vultures (*Coragyps atratus*; 0.6788), unspecified *Falconidae* (0.5556), and rough-legged hawks (*Buteo lagopus*; .05000). Of the remaining *Cathartidae*, prevalence rates in turkey vultures (*Cathartes aura*; 0.3925) and unspecified *Cathartidae* (0.3529) were higher than that of California condors (*Vultur gryphus*; 0.0375). In the *Accipitridae* family, prevalence was highest in red-tailed hawks (0.2584), followed by bald eagles (0.2557), Swainson’s hawks (*Buteo swainsoni*; 0.2105), unspecified eagles (0.1538), unspecified hawks (0.1282), and red-shouldered hawks (*Buteo lineatus*; 0.1117). The prevalence rates of the remaining *Accipitridae* species tested were below 0.1000. Following the prairie falcon and unspecified *Falconidae*, peregrine falcons had a prevalence of 0.3108. The remaining *Falconidae* species had prevalence rates below 0.1000. Osprey (*Pandion haliaetus*), the only species within *Pandionidae,* had a prevalence rate of 0.0488. Prevalence in the *Strigidae* family was highest in great horned owls (0.3836), followed by short-eared owls (*Asio flammeus*; 0.3333) and snowy owls (*Bubo scandiacus*; 0.3214). Barred owls (*Strix varia)*, eastern screech-owls (*Megascops asio*), long-eared owls (*Asio otus*), and unidentified *Strigidae* all had prevalence rates below 0.1000.

**Table 5.** Avian influenza A virus detections in Galliformes morbidity/mortality submissions as reported by diagnostic laboratories in the NAHLN from 1 February 2022 to 31 March 2023.

| Family | Species | Number of Birds Sampled | Influenza Type A rRT-PCR Detections (N Positive) | Prevalence |
| --- | --- | --- | --- | --- |
|  | Greater sage grouse | 5 | 1 | 0.2000 |
|  | Pheasant (unidentified) | 674 | 61 | 0.0905 |
|  | Ring-necked pheasant | 233 | 31 | 0.1330 |
|  | Ruffed grouse | 15 | 1 | 0.0667 |
|  | Wild turkey | 451 | 29 | 0.0643 |
|  | Other *Phasianidae* spp. | 52 | 0 | 0 |
| *Phasianidae* |  | 1430 | 123 | 0.0860 |
| *Odontophoridae* | Quail (unidentified) | 757 | 3 | 0.0040 |
| Total |  | 3617 | 249 | 0.0688 |

**Table 6.** Avian influenza A virus detections in raptor morbidity/mortality submissions as reported by diagnostic laboratories in the NAHLN from 1 February 2022 to 31 March 2023.

| Family | Species | Number of Birds Sampled | Influenza Type A rRT-PCR Detections (N Positive) | Prevalence |
| --- | --- | --- | --- | --- |
|  | Bald eagle | 1150 | 294 | 0.2557 |
|  | Broad-winged hawk | 120 | 6 | 0.0500 |
|  | Coopers hawk | 279 | 22 | 0.0789 |
|  | Golden eagle | 69 | 3 | 0.0435 |
|  | Red-shouldered hawk | 179 | 20 | 0.1117 |
|  | Red-tailed hawk | 747 | 193 | 0.2584 |
|  | Rough-legged hawk | 16 | 8 | 0.5000 |
|  | Sharp-shinned hawk | 55 | 4 | 0.0727 |
|  | Swainson's hawk | 19 | 4 | 0.2105 |
|  | Hawk (unidentified) | 78 | 10 | 0.1282 |
|  | Eagle (unidentified) | 13 | 2 | 0.1538 |
|  | Other *Accipitridae* spp. | 43 | 0 | 0 |
| *Accipitridae* |  | 2768 | 566 | 0.2044 |
|  | Black vulture | 495 | 336 | 0.6788 |
|  | California condor | 80 | 3 | 0.0375 |
|  | Turkey vulture | 186 | 73 | 0.3925 |
|  | *Cathartidae* (unidentified) | 34 | 12 | 0.3529 |
| *Cathartidae* |  | 795 | 424 | 0.5333 |
|  | American kestrel | 101 | 3 | 0.0297 |
|  | Merlin | 46 | 4 | 0.0870 |
|  | Peregrine falcon | 148 | 46 | 0.3108 |
|  | Prairie falcon | 1 | 1 | 1.0000 |
|  | *Falconidae* (unidentified) | 9 | 5 | 0.5556 |
|  | Other *Falconidae* spp. | 82 | 0 | 0 |
| *Falconidae* |  | 387 | 59 | 0.1525 |
| *Pandionidae* | Osprey | 82 | 4 | 0.0488 |
|  | Barred owl | 320 | 23 | 0.0719 |
|  | Eastern screech-owl | 96 | 3 | 0.0313 |
|  | Great horned owl | 610 | 234 | 0.3836 |
|  | Long-eared owl | 11 | 1 | 0.0909 |
|  | Short eared owl | 3 | 1 | 0.3333 |
|  | Snowy owl | 28 | 9 | 0.3214 |
|  | *Strigidae* (unidentified) | 108 | 10 | 0.0926 |
|  | Other *Strigidae* spp. | 36 | 0 | 0 |
| *Strigidae* |  | 1213 | 281 | 0.2317 |
| *Tytonidae* | Barn owl | 61 | 0 | 0 |
| Total |  | 5306 | 1334 | 0.2514 |

### Pheasants, Turkeys, and Quail: Order Galliformes

Out of the 3,617 Galliformes species submitted for testing, 249 tested positive for AI viruses, resulting in a total prevalence of 0.0688 (Table 5). Prevalence rates within the *Phasianidae and Odontophoridae* families were 0.0860 and 0.0040, respectively. Of the species tested within *Odontophoridae*, prevalence was highest in greater sage grouse (*Centrocercus urophasinus;* 0.2000), followed by ring-necked pheasant (*Phasianus colchicus;* 0.1330), unspecified pheasants (0.0905), ruffled grouse (*Bonasa umbellus;* 0.0667), and wild turkey (*Meleagris gallopavo; 0.0643*).

### National Veterinary Services Laboratories

A total of 2,121 samples from our synanthropic species of interest were confirmed as the Eurasian lineage Gs/GD H5 clade 2.3.4.4b subtype at the NVSL between 1 February 2022 and 31 March 2023 (Table 7). Detections from orders Anseriformes, Charadriiformes, Pelecaniformes, Suliformes, and Gruiformes were excluded from our dataset. Of the remaining orders, approximately 92% of the samples (1,942) originated from raptors: 840 Accipitriformes, 671 Cathartiformes, 61 Falconiformes, and 370 Strigiformes. Detections also were confirmed in 149 Passeriformes and 30 Galliformes.

**Table 7.** Prevalence in avian orders with confirmed HPAI EA H5 detections as determined by rRT-PCR assay targeting Eurasian lineage Gs/GD H5 clade 2.3.4.4b at the NVSL from 1 February 2022 to 31 March 2023.

| Order | H5 2.3.4.4b rRT-PCR Detections |
| --- | --- |
| Accipitriformes | 840 |
| Cathartiformes | 671 |
| Falconiformes | 61 |
| Galliformes | 30 |
| Passeriformes | 149 |
| Strigiformes | 370 |
| Total | 2,121 |

## Discussion

### Targeted Surveillance

Based on rRT-PCR results, we did not detect any AI viruses (HPAI or other) in the 266 wild birds we sampled at two commercial poultry premises with confirmed poultry detections of H5N1 in Dubois County, IN. A total of three commercial poultry premises in Dubois County were confirmed positive for H5N1 during February 2022, and anecdotal reports confirm flocks of migrant European starlings and mixed blackbird species in the area. It is possible that the virus was present in wild bird species around these premises; however, factors in our sampling methods may have negatively impacted the ability to detect AI viruses. First, surveillance began after the commercial facilities were quarantined and poultry were euthanized, potentially preventing the capture of wild birds that may have been utilizing poultry barns. Further, the approximate two-week delay between H5N1 confirmation at the premises and the initiation of wild bird surveillance might have contributed to the lack of detections. Other studies similarly noted that such a delay may have contributed to a lack of HPAI virus detections [18,47]. Second, our study did not investigate non-infected farms, but sampling at non-infected farms in conjunction with infected farms could provide a more comprehensive view of disease ecology and host population dynamics in the area [48]. Third, our low sample size, approximately 130 birds per farm, may have influenced the ability to detect any AI viruses in captured species.

Similar limitations in the surveillance of synanthropic birds on HPAI infected farms have been noted in previous investigations [18]. Enhanced surveillance with a sufficient sample size of wild birds in known areas of HPAI virus detections in poultry is essential to understand disease ecology and the role potential bridge hosts play in transmission [49,50]. Conducting future sampling concurrent with poultry depopulation activities, minimizing the delay between the confirmation of HPAI and initiation of wild bird sampling, and investigating populations at uninfected farms all could provide a more comprehensive picture of wild bird – poultry transmission risk and directionality.

While this investigation suggests that synanthropic species minimally contribute to the spread of HPAI to poultry, there are inherent limiting factors that may have underrated the perceived risk of transmission. Synanthropic birds may die quickly once infected and their probability of capture is lower than that of healthy individuals, resulting in a potential underestimation of disease prevalence [18,47]. Further, as passerine species tend to be smaller in size than raptors or waterfowl species, moribund passerines may have a lower detection probability due to a smaller distribution of feathers and bones or quick removal by scavengers or predators [49,51]. Wobeser and Wobeser [52] found approximately 70% of small bird carcasses experimentally placed were removed within 24 hours by natural means and noted the presence of several scavenging species during that time frame. Although rates of carcass removal are site specific and variable, evidence indicates the probability of detecting a species is negatively correlated with both the length of time post mortality and the size of the birds.

Full length viral genome sequence analyses of 1,369 HPAI H5N1 detections in wild birds, commercial poultry, and backyard flocks from December 2021 to April 2022, suggest that at least 85% of U.S. HPAI virus detections in poultry premises and non-poultry flocks are consistent with wild bird origin, while approximately 15% of detections are consistent with lateral transmission (poultry to poultry) [15]. This suggests that wild birds are major contributors to the spread of HPAI H5N1 to poultry, and environmental contamination or direct transmission from a variety of wild bird species are potential sources. Further research is needed to understand the transmission pathways from wild birds to poultry.

Conducting risk assessments and determining wild bird activity on farms can be used to increase biosecurity and protect domestic poultry populations [53]. Knowledge of the wild bird – poultry interface, species of concern, and the space where interspecific interactions occur is critical in developing biosecurity methods to decrease contact and risk of AI virus transmission [31].

Understanding the disease ecology and risk of viral transmission could aid producers in minimizing the risk to poultry by reducing attractants and contact between wild birds and poultry on farms. Although AI viruses previously have been detected experimentally in passerine species, including five out of the eight species sampled during targeted surveillance, both targeted sampling and M/M investigations throughout the ongoing 2022 – 2023 H5N1 outbreak in the USA show low prevalence in this order [2]. More research is needed to determine which wild bird species may be involved in viral transmission to domestic poultry.

### Morbidity/Mortality Investigations

The 2022 – 2023 outbreak of HPAI H5N1 was widespread in wild avifauna, with virus detections across the conterminous U.S. and Alaska in synanthropic orders Accipitriformes, Cathartiformes, Falconiformes, Galliformes, Passeriformes, and Strigiformes. Prevalence rates of AI virus detections from 1,666 M/M samples from 1 February 2022 to 31 March 2023, tested at the NAHLN were highest in vultures (0.5333) followed by owls (0.2318), eagles and hawks (0.2044), falcons (0.1525), corvids (0.1240), pheasants and grouse (0.0860), songbirds (0.0147), doves (0.0090), and quail (0.0040). Confirmatory testing by the NVSL of over 2,100 samples across the same orders and timeframe suggests that HPAI H5N1 was the predominant strain circulating and causing morbidity and mortality in wild bird populations in the USA.

Avian ecology and behavior likely play a major role in the transmission of the virus. Predatory and scavenging species show substantially increased levels of infection when compared to granivorous or insectivorous groups, suggesting that transmission may occur via consumption of infected birds or mammals [40]. The order Accipitriformes had the greatest disease prevalence overall, of which vultures, the only obligate scavenger sampled, had the highest rate of infection. Furthermore, roosting behavior, such as displayed in vulture species, increases sociality between conspecifics and the likelihood of viral transmission, particularly for density-dependent pathogens such as AI viruses that spread fecal – orally [54,55]. Facultative scavenging raptors, such as hawks, eagles, owls, and falcons, consume both carrion and apparently healthy prey, which may explain the lower prevalence rates in these families. Previous research of HPAI susceptibility in raptor species supports these findings. Uno et al. [56] found high levels of HPAI H5N1 infection in kestrels following experimental inoculation or ingestion of infected poultry meat. Furthermore, captive raptor morbidities and mortalities during the 2014 – 2015 outbreak were attributed to ingestion of infected meat [13]. Investigating families based on diet may help explain why *Corvidae*, with frequent scavenging behavior and a higher probability of feeding upon infected animals [57], have a prevalence of 0.1240 compared to approximately 0.0147 in non-omnivorous songbirds. It is possible that ingestion of infected tissue is a key transmission pathway from scavenging species to conspecifics, heterospecifics, or domestic poultry.

Although there has been previous concern about high potential rates of infection in Galliformes due to their close association with humans and domestic poultry [58], our observed rates of infection are only slightly higher in Galliformes (0.0688) than songbirds (0.0147) and Columbiformes (0.0090). These groups have similar diets, ecological niches, and contact rates with conspecifics, humans, and domestic animals [58], suggesting that factors influencing AI transmission may go beyond physiology and behavior. Non-predatory species tend to have increased sociality [59]. Thus, lower prevalence rates in these groups suggest that the risk of transmission by direct contact with conspecifics is low. However, as virus was detected in these groups, alternative transmission pathways beyond oral consumption and contact with conspecifics should be considered. While experimental research has shown the potential for AI viruses to be transmitted between species via shared environmental resources such as water sources [60,61], further investigation is needed to understand AI virus transmission across the landscape in free ranging avian populations.

Understanding AI virus transmission is critical to protect and manage wild bird populations, especially threatened and endangered species. Raptor species, particularly those with smaller population sizes and geographical ranges (e.g., California condors [*Gymnogyps californianus*]), that scavenge or prey upon other avian species have a higher risk of deleterious population impacts caused by HPAI virus infections [56]. Bertran et al. [39] note that the introduction of HPAI viruses in raptors could negatively impact already threatened species and surveillance may be an invaluable tool to better understand the epidemiology of AI viruses in these populations.

An understanding of the increased risk for scavenging species has already been applied to management strategies meant to protect the highly endangered California condor, including vaccination and increased surveillance efforts [62]. Monitoring sensitive species (e.g., conducting active surveillance or risk assessments) during an HPAI outbreak can offer valuable information to wildlife managers on population dynamics, disease risk, and virus type and distribution. Identifying susceptible species with fragile populations could aid in conservation efforts.

Sampling birds as part of M/M investigations may have introduced bias into the dataset as it is more probable to detect disease in these groups than in apparently healthy birds. Further, more charismatic species such as raptors may have had disproportionate detections due to birds being larger, more noticeable, and more publicly valued. However, this methodology allowed for the largest possible dataset, potentially increasing the precision of estimates. The 67% response rate from labs within the NAHLN and the differences in each lab’s Laboratory Information Management System taxonomy lists may have restricted the ability to draw comprehensive conclusions on AI virus ecology in different avian orders. Expanding future investigations to include apparently healthy wildlife in conjunction with M/M investigations could provide key insights into the disease ecology of AI viruses and their implications for wildlife, human, and agricultural health.

## Conclusions

Active surveillance of wild birds at HPAI infected poultry facilities combined with morbidity and mortality surveillance of synanthropic birds offers an avenue to better understand the ecology of avian influenza viruses and the risks they pose to wildlife, domestic animals, and human health. No virus was detected through active surveillance in the orders Columbiformes and Passeriformes. Further, the lowest prevalence rates from morbidity and mortality investigations were observed in Columbiformes, Passeriformes, and Galliformes. Our results suggest that these orders pose a lower risk of acting as major transmission pathways of AI viruses compared to the orders Cathartiformes, Strigiformes, Accipitriformes, and Falconiformes. The most prevalent viral detections were found in wild predatory, scavenging birds, suggesting that there is strong evidence that the consumption of infected tissue is a key pathway for the transmission of AI viruses in these species. Understanding the factors influencing AI virus transmission is crucial for the development and implementation of superior management strategies.

## Data Availability

Data for highly pathogenic avian influenza detections in wild birds confirmed at the NVSL from 2022 to 2023 are available at <u>USDA APHIS | 2022-2023 Detections of Highly Pathogenic Avian Influenza in Wild Birds</u>. The majority of data supporting this research are restricted and not available publicly. Wild bird influenza surveillance data collected between August 2007 and July 2023 are available from the Wildlife Services National Wildlife Disease Program (NWDP) of the USDA by contacting the NWDP at.

## Acknowledgements

We are very appreciative of the cooperation and support of the poultry farms for allowing us access to their properties and for on-site support. We thank the IN Board of Animal Health, the IN Department of Natural Resources – Division of Fish and Wildlife, the IN Division of Law Enforcement, the IL Department of Natural Resources, the IN Department of Agriculture, the IN Division of the USFWS, and Veterinary Services for their expertise and assistance in planning and wild bird sampling. We thank the NAHLN and the NVSL for their support and allowing access to their data. Finally, we thank the IL and IN Wildlife Services field operation personnel for their hard work to make this project successful.

## Supplementary Materials

Compiled dataset of all families, species, number of birds sampled, and positive detections as determined by a general influenza Type A rRT-PCR assay across all samples submitted as part of morbidity/mortality events that were tested at diagnostic laboratories in the NAHLN from 1 February 2022 to 31 March 2023.

## References

[1] Canadian Food Inspection Agency [CFIA]. (2023, June 07). Status of ongoing avian influenza response by province. Retrieved June 08, 2023, from https://inspection.canada.ca/animal-health/terrestrial-animals/diseases/reportable/avian-influenza/HPAIV-in-canada/status-of-ongoing-avian-influenza-response/eng/1640207916497/1640207916934

[2] U.S. Department of Agriculture [USDA], Animal and Plant Health Inspection Service. (2023a). 2022-2023 detections of highly pathogenic avian influenza in wild birds [Data set]. Retrieved from https://www.aphis.usda.gov/aphis/ourfocus/animalhealth/animal-disease-information/avian/avian-influenza/hpai-2022/2022-hpai-wild-birds

[3] Zhang, G., Li, B., Raghwani, J., Vrancken, B., Jia, R., Hill, S. C.,…Tian, H. (2023). Bidirectional movement of emerging H5N8 avian influenza viruses between Europe and Asia via migratory birds since early 2020. Molecular Biology and Evolution, 40(2). 10.1093/molbev/msad019

[4] Harvey, J. A., Mullinax, J. M., Runge, M. C., & Prosser, D. J. (2023). The changing dynamics of highly pathogenic avian influenza H5N1: next steps for management & science in North America. Biological Conservation, 283. 10.1016/j.biocon.2023.110041

[5] Bevins, S. N., Shriner, S. A., Cumbee, J. C., Dilione, K. E., Douglass, K. E., Ellis, J. W.,…Lenoch, J. B. (2022). Intercontinental movement of highly pathogenic avian influenza A (H5N1) clade 2.3.4.4 virus to the United States, 2021. Emerging Infectious Diseases, 28(5), 1006–1011. 10.3201/eid2805.220318

[6] Fouchier, R. A. M., Munster, V., Wallensten, A., Bestebroer, T. M., Herfst, S., Smith, D.,…Osterhaus, A. D. M. E. (2005). Characterization of a novel influenza A virus hemagglutinin subtype (H16) Obtained from Black-Headed Gulls. Journal of Virology, 79(5), 2814–2822. 10.1128/JVI.79.5.2814-2822.2005

[7] Gauthier-Clerc, M., Lebarbenchon, C., & Thomas, F. (2007). Recent expansion of highly pathogenic avian influenza H5N1: a critical review. IBIS, 149(2), 202–214. 10.1111/j.1474-919X.2007.00699.x

[8] Olsen, B., Munster, V. J., Wallensten, A., Walenström, J., Osterhaus, A. D. M. E., & Fouchier, R. A. M. (2006). Global patterns of influenza A virus in wild birds. Science, 312(5772). 10.1126/science.1122438

[9] Ramey, A. M., Hill, N. J., DeLiberto, T. J., Gibbs, S. E. J., Camille Hopkins, M., Lang, A. S.,…Wan, X. (2022). Highly pathogenic avian influenza is an emerging disease threat to wild birds in North America. The Journal of Wildlife Management, 86(2). 10.1002/jwmg.22171

[10] Ip, H. S., Dusek, R. J., Bodenstein, B., Torchetti, M. K., DeBruyn, P., Mansfield, K. G.,…Sleeman, J. M. (2016). High rates of detection of clade 2.3.4.4 highly pathogenic avian influenza H5 viruses in wild birds in the Pacific Northwest during the winter of 2014-15. Avian Disease, 60, 354-358. 10.1637/11137-050815-Reg

[11] Suarez, D. L. (2017). Influenza A virus. In Animal Influenza (pp. 3-30). Retrieved from https://onlinelibrary.wiley.com/templates/jsp/_ux3/_acropolis/_pericles/pdf-viewer/web/viewer.html?file=/doi/pdfdirect/10.1002/9781118924341

[12] Yoon, S-W., Webby, R. J., & Webster, R. G. (2014). Evolution and ecology of influenza A viruses. In Influenza Pathogenesis and Control (pp. 359-375). Retrieved from https://link.springer.com/chapter/10.1007/82_2014_396

[13] U.S. Department of Agriculture [USDA], Animal and Plant Health Inspection Service. (2016). Final report for the 2014-2015 outbreak of highly pathogenic avian influenza (HPAI) in the United States. Retrieved from https://www.aphis.usda.gov/animal_health/emergency_management/downloads/hpai/2015-hpai-final-report.pdf

[14] U.S. Department of Agriculture [USDA], Animal and Plant Health Inspection Service. (2023b). 2022-2023 detections of highly pathogenic avian influenza in commercial and backyard flocks [Data set]. Retrieved from https://www.aphis.usda.gov/aphis/ourfocus/animalhealth/animal-disease-information/avian/avian-influenza/hpai-2022/2022-hpai-commercial-backyard-flocks

[15] Youk, S., Torchetti, M. K., Lantz, K., Lenoch, J. B., Killian, M. L., Leyson, C.,…Pantin-Jackwood, M. J. (2023). H5N1 highly pathogenic avian influenza clade 2.3.4.4b in wild and domestic birds: introductions into the United States and reassortments, December 2021-April 2022. Virology, 587. 10.1016/j.virol.2023.109860

[16] Ssematimba, A., Hagenaars, T. J., de Wit, J. J., Ruiterkamp, F., Fabri, T. H., Stegeman, J. A., & de Jong, M. C. M. (2013). Avian influenza transmission risks: analysis of biosecurity measures and contact structure in Dutch poultry farming. Preventative Veterinary Medicine, 109, 106–115. 10.1016/j.prevetmed.2012.09.001

[17] Caron, A., Cappelle, J., Cumming. G.S., de Garine-Wichatitsky, M., & Gaidet, N. (2015). Bridge hosts, a missing link for disease ecology in multi-host systems. Veterinary Research, 46(83). doi: 10.1186/s13567-015-0217-9

[18] Shriner, S. A., Root, J. J., Lutman, M. W., Kloft, J. M., VanDalen, K. K., Sullivan, H. J.,…DeLiberto, T. J. (2016). Surveillance for highly pathogenic H5 avian influenza virus in synanthropic wildlife associated with poultry farms during an acute outbreak. Scientific Reports, 6(1), 36237. 10.1038/srep36237

[19] Shriner, S. A., & Root, J. J. (2020). A review of avian influenza A virus associations in synanthropic birds. Viruses, 12(11), 1209. 10.3390/v12111209

[20] Abolnik, C. (2014). A current review of avian influenza in pigeons and doves (Columbidae). Veterinary Microbiology, 170(3-4), 181–196. 10.1016/j.vetmic.2014.02.042

[21] Brown, J. D., Stallknecht, D. E., Berghaus, R. D., & Swayne, D. E. (2009). Infectious and lethal doses of H5N1 highly pathogenic avian influenza virus for house sparrows (*Passer Domesticus*) and rock pigeons (*Columbia Livia*). Journal of Veterinary Diagnostic Investigation, 21(4), 437–445. 10.1177/104063870902100404

[22] Caron, A., Cappelle, J., & Gaidet, N. (2017). Challenging the conceptual framework of maintenance hosts for influenza A viruses in wild birds. Journal of Applied Ecology, 54, 681–690. 10.1111/1365-2664.12839

[23] Kwon, Y-K., Joh, S-J., Kim, M-C., Lee, Y-J., Choi, J-G., Lee, E-K.,…Kim, J-H. (2005). Highly pathogenic avian influenza in magpies (*Pica pica sericea*) in South Korea. Journal of Wildlife Diseases, 41(3), 618–623. 10.7589/0090-3558-41.3.618

[24] Tanimura, N., Tsukamoto, K., Okamatsu, M., Mse, M., Imada, T., Nakamura, K.,…Imai, K. (2006). Pathology of fatal highly pathogenic H5N1 avian influenza virus infection in large-billed crows (*Corvus macrorhynchos*) during the 2004 outbreak in Japan. Veterinary Pathology, 43(4), 500–509. 10.1354/vp.43-4-500

[25] Kumar, M., Murugkar, H.V., Nagarajan, S., Tosh, C., Patil, S., Nagaraja, K. H.,…Dubey, S. C. (2020). Experimental infection and pathology of two highly pathogenic avian influenza H5N1 viruses isolated from crow and chicken in hous crows (Corvus splendens). Acta Virologica, 64, 325–330. 10.4149/av_2020_306

[26] Soda, K., Tomioka, Y., Usui, T., Ozaki, H., Yamaguchi, T., & Ito, T. (2020). Pathogenicity of H5 highly pathogenic avian influenza virus in rooks (Corvus frugilegus). Avian Pathology, 49(3), 261–267. 10.1080/03079457.2020.1724876

[27] Biswas, M., Rahman, M. H., Das, A., Ahmed, S. S. U., Giasuddin, M., Christensen, J. P. (2011). Risk for highly pathogenic avian influenza H5N1 virus infection in chickens in small-scale commercial farms, in a high-risk area, Bangladesh, 2008. Transboundary and Emerging Diseases, 58, 519-525. 10.1111/j.1865-1682.2011.01235.x

[28] Caron, A., Chiweshe, N., Mundava, J., Abolnik, C., Capobianco Dondona, A., Scacchia, M., & Gaidet, N. (2017). Avian viral pathogens in swallows, Zimbabwe: infectious diseases in Hirundinidae: a risk to swallow? Ecohealth, 14(4), 805-809.

[29] Gronesova, P., Kabat, P., Trnka, A., & Betakova, T. (2008). Using nested RT-PCR analyses to determine the prevalence of avian influenza viruses in passerines in western Slovakia, during summer 2007. Scandinavian Journal of Infectious Diseases, 40, 954–957. 10.1080/00365540802400576

[30] Caron, A., Grosbois, V., Etter, E., Gaidet, N., de Garine-Wichatitsky, M. (2014). Bridge hosts for avian influenza viruses at the wildlife/domestic interface: an eco-epidemiological framework implemented in southern Africa. Preventive Veterinary Medicine, 117(2-4), 590–600. 10.1016/j.prevetmed.2014.09.014

[31] Valdez-Gómez, H. E., & Hernandez, I. G. (2017). Risk factors for the transmission of infectious diseases agents at the wild birds-commercial birds interface. A pilot study in the region of the Altos de Jalisco, Mexico. Bulletin de l’Acadamie Veterinaire de France, 170(2), 143–150. 10.4267/2042/62332

[32] Hiono, T., Okamatsu, M., Yamamoto, N., Ogasawara, K., Endo, M., Kuribayashi, S.,…Sakoda, Y. (2016). Experimental infection of highly and low pathogenic avian influenza viruses to chickens, ducks, tree sparrows, jungle crows, and black rats for the evaluation of their roles in virus transmission. Veterinary Microbiology, 182, 108–115. 10.1016/j.vetmic.2015.11.009

[33] Ellis, J. W., Root, J. J., McCurdy, L. M., Bentler, K. T., Barrett, N. L., Van Dalen, K. K.,…Shriner, S. A. (2021). Avian influenza A virus susceptibility, infection, transmission, and antibody kinetics in European starlings. PLOS Pathogens, 17(1). 10.1371/journal.ppat.1009879

[34] Root, J. J., Ellis, J. W., & Shriner, S. A. (2022). Strength in numbers: avian influenza A virus transmission to poultry from a flocking passerine. Transboundary and Emerging Diseases, 69, 1153–1159. 10.1111/tbed.14397

[35] Fuller, T. L., Saatchi, S. S., Curd, E. E., Toffelmier, E., Thomassen, H. A., Buermann, W.,…Smith, T. B. (2010). Mapping the risk of avian influenza in wild birds in US. BMC Infectious Diseases, 10(187). 10.1186/1471-2334-10-187

[36] Root, J. J., Bosco-Lauth, A. M., Marlenee, N. L., & Bowen, R. A. (2018). Viral shedding of clade 2.3.4.4 H5 highly pathogenic avian influenza viruses by American robins. Transboundary and Emerging Diseases, 65(6), 1823-1827. 10.1111/tbed.12959

[37] De Marco, M.A., Campitelli, L., Delogu, M., Raffini, E., Foni, E., Di Trani, L.,…Donatelli, I. (2005). Serological evidences showing the involvement of free-living pheasants in the influenza ecology. Animal Science, 4, 287–291. doi: 10.4081/ijas.2005.287

[38] Ferro, P. J., Khan, O., Vuong, C., Reddy, S. M., LaCoste, L., Rollins, D., & Lupiani, B. (2012). Avian influenza virus investigation in wild bobwhite quail in Texas. Avian Diseases, 56, 858–860. 10.1637/10197-041012-ResNote.1

[39] Bertran, K., Busquets, N., Xavier A., Francesca, Garcia de la Fuete J., (third), Solanes, D.,…Majo, N., (second). (2012). Highly (H5N1) and low (h7N2) pathogenic avian influenza virus infection in falcons via nasochoanal route and ingestion of experimentally infected prey. PLoS One, 7(3). 10.1371/journal.pone.0032107

[40] Hall, J. S., Ip, H. S., Franson, J. C., Meteyer, C., Nashold, S., TeSlaa, J. L., French, J.,…Brand, C. (2009). Experimental infection of a North American raptor, American kestrel (Falco sparverius), with highly pathogenic avian influenza virus (H5N1). PLoS ONE, 4(10). 10.1371/journal.pone.0007555

[41] Kocan, A. A., Snelling, J., & Greiner, E. C. (1977). Some infectious 10.7589/0090-3558-13.3.304 and parasitic disease in Oklahoma raptors. Journal of Wildlife Diseases, 13, 304-306.

[42] Redig, P. T. & Goyal, S. M. (2012). Serologic evidence of exposure of raptors to influenza a virus. Avian Diseases, 56, 411–413. 10.1637/9909-083111-ResNote.1

[43] Goyal, S. M., Jindal, N., Chander, Y., Ramakrishnan, M. A., Redig, P. T., & Sreevatsan, S. (2010). Virology Journal, 7. 10.1186/1743-422X-7-174

[44] Manvell, R. J., McKinney, P., Wernery, U., & Frost, K. (2000). Isolation of a highly pathogenic influenza a virus subtype H7N3 from a peregrine falcon. Isolation of a highly pathogenic influenza a virus subtype H7N3 from a peregrine falcon. Avian Pathology, 29, 635-637. 10.1080/03079450020016896

[45] Spackman, E., Senne, D. A., Myers, T. J., Bulaga, L. L., Garber, L. P., Perdue, M. L.,…Suarex, D. L. (2002). Development of real-time reverse transcriptase PCR assay for type A influenza virus and the avian H5 and H7 hemagglutinin subtypes. Journal of Clinical Microbiology, 40(9), 3256–3260. doi: 10.1128/JCM.40.9.3256-3260.2002

[46] The National Animal Health Laboratory Network [NAHLN] Standard Operating Procedure for Real-time RT-PCR Detection of Influenza A and Avian Paramyxovirus Type-1 (NVSL-SOP-0068).

[47] Grear, D. A., Dusek, R. J., Walsh, D. P., & Hall, J. S. (2017). No evidence of infection or exposure to highly pathogenic avian influenzas in peridomestic wildlife on an affected poultry facility. Journal of Wildlife Diseases, 53(1), 37. 10.7589/2016-02-029

[48] Hoye, B. J., Munster, V. J., Nishiura, H., Klaasman, M., & Fouchier, R. A. M. (2010). Surveillance of wild birds for avian influenza virus. Emerging Infectious Diseases, 16(12), 1827–1834.

[49] Hesterberg, U., Harris, K., Stroud, D., Guberti, V., Busani, L., Pittman, M.,…Brown, I. (2009). Avian influenza surveillance in wild birds in the European Union in 2006. Influenza and Other Respiratory Viruses, 3(1), 1–14. 10.1111/j.1750-2659.2008.00058.x

[50] Ward, M. P. (2007). Geographic information system-based avian influenza surveillance systems for village poultry in Romania. Veterinaria Italiana, 43(3), 483–489.

[51] Uddin, M., Dutta, S., Kolipakam, V., Sharma, H., Usmani, F., & Jhala, Y. (2021). High bird mortality due to power lines invokes urgent environmental mitigation in a tropical desert. Biological Conservation, 261. 10.1016/j.biocon.2021.109262

[52] Wobeser, G. & Wobeser, A. G. (1992). Carcass disappearance and estimation of mortality in a simulated die-off of small birds. Journal of Wildlife Disease, 28(4), 548–554. 10.7589/0090-3558-28.4.548

[53] Burns, T. E., Ribble, C., Stephen, C., Kelton, D., Toews, L., Osterhold, J., & Wheeler, H. (2012). Use of observed wild bird activity on poultry farms and a literature review to target species as high priority for avian influenza testing in 2 regions of Canada. The Canadian Veterinary Journal = La Revue Veterinaire Canadienne, 53(2), 158–166. Retrieved from https://www.canadianveterinarians.net/journals-and-classified-ads/the-canadian-veterinary-journal/

[54] Avery, M. L, & Lowney, M. S. (2016, October). Vultures. In Wildlife Damage Management Technical Series. Retrieved from https://digitalcommons.unl.edu/cgi/viewcontent.cgi?article=1004&context=nwrcwdmts

[55] Laughlin, A. J., Hall, R. J., & Taylor, C. M. (2019). Ecological determinants of pathogen transmission in communally roosting species. Theoretical Ecology, 12, 225–235. 10.1007/s12080-019-0423-6

[56] Uno, Y., Soda, K., Tomioka, Y., Ito, T., Usui, T., & Yamaguchi, T. (2020). Pathogenicity of clade 2.3.2.1 H5N1 highly pathogenic avian influenza virus in American kestrel (Falco sparverius). Avian Pathology, 49(5), 515-525. 10.1080/03079457.2020.1787337

[57] Bragato, P. J., Spencer, E. E., Dickman, C. R., Crowther, M. S., Tulloch, A., & Newsome, T. M. (2022). Effects of habitat, season and flood on corvid scavenging dynamics in Central Australia. Austral Ecology, 47, 939–953. 10.1111/aec.13177

[58] Crespo, R., Franca, M. S., Fenton, H., & Shivaprasad, H. L. (2018). Galliformes and Columbiformes. In Pathology of Wildlife and Zoo Animals (pp. 747-773). Retrieved from https://www.sciencedirect.com/science/article/pii/B9780128053065000316

[59] Song, Z., Liker, A., Liu, Y., & Szekely, T. (2022). Evolution of social organization: phylogenetic analyses of ecology and sexual selection in weavers. The American Naturalist, 200(2), 181–301. 10.1086/720270

[60] Root, J. J., Shriner, S. A., Ellis, J. W., Vandalen, K. K., & Sullivan, H. J. (2015). When fur and feather occur together: interclass transmission of avian influenza A virus from mammals to birds through common resources. Scientific Reports, 5. 10.1038/srep14354

[61] VanDalen, K. K., Franklin, A. B., Mooers, N. L., Sullivan, H. J., & Shriner, S. A. (2010). Shedding light on avian influenza H4N6 infection in mallards: modes of transmission and implications for surveillance. PLoS One, 5(9).

[62] U.S. Fish and Wildlife Service [USFWS]. (2023). HPAI update. In California Condor Recovery Program. Retrieved from https://www.fws.gov/program/california-condor-recovery/southwest-california-condor-flock-hpai-information-updates-2023

