## Supplemental Table 1 for "Prevalence of Avian Influenza Virus in Synanthropic Birds Associated with an Outbreak of Highly Pathogenic Strain EA/AM H5N1"

### Supplementary Material

Table S1. Families, species, number of birds sampled, and positive detections as determined by a general influenza Type A rRT-PCR assay across all samples submitted as part of morbidity/mortality events that were tested at diagnostic laboratories in the NAHLN from 1 February 2022 to 31 March 2023.

| Family | Common Name | Scientific Name | Number of Birds Sampled | Virus/RNA Detection (N Positive) |
|---|---|---|---|---|
| *Columbidae* | Rock dove | *Columba livia* | 244 | 2 |
| *Columbidae* | Common ground dove | *Columbina passerina* | 4 | 0 |
| *Columbidae* | Eurasian collared dove | *Streptopelia decaocto* | 11 | 0 |
| *Columbidae* | White-winged dove | *Zenaida asiatica* | 3 | 0 |
| *Columbidae* | Mourning dove | *Zenaida macroura* | 92 | 2 |
| *Columbidae* | Dove (unidentified) | | 54 | 0 |
| *Columbidae* | Pigeon (unidentified) | | 35 | 0 |
| *Bombycillidae* | Cedar waxwing | *Bombycilla cedrorum* | 20 | 0 |
| *Bombycillidae* | Bohemian waxwing | *Bombycilla garrulus* | 1 | 0 |
| *Cardinalidae* | Northern cardinal | *Cardinalis cardinalis* | 9 | 0 |
| *Cardinalidae* | Indigo bunting | *Passerina cyanea* | 3 | 0 |
| *Cardinalidae* | Rose-breasted grosbeak | *Pheucticus ludovicianus* | 2 | 0 |
| *Cardinalidae* | Scarlet tanager | *Piranga olivacea* | 2 | 0 |
| *Fringillidae* | Common redpoll | *Acanthis flammea* | 2 | 0 |
| *Fringillidae* | House finch | *Haemorhous mexicanus* | 10 | 0 |
| *Fringillidae* | Evening grosbeak | *Hesperiphona vespertina* | 1 | 0 |
| *Fringillidae* | Pine grosbeak | *Pinicola enucleator* | 4 | 1 |
| *Fringillidae* | American goldfinch | *Spinus tristis* | 19 | 1 |
| *Fringillidae* | Finch (unidentified) | | 11 | 0 |
| *Hirundinidae* | Barn swallow | *Hirundo rustica* | 12 | 0 |
| *Hirundinidae* | Tree swallow | *Tachycineta bicolor* | 19 | 2 |
| *Hirundinidae* | Violet-green swallow | *Tachycineta thalassina* | 4 | 3 |
| *Icteridae* | Red-winged blackbird | *Agelaius phoeniceus* | 9 | 1 |
| *Icteridae* | Bobolink | *Dolichonyx oryzivorus* | 1 | 0 |
| *Icteridae* | Brown-headed cowbird | *Molothrus ater* | 7 | 0 |
| *Icteridae* | Boat-tailed grackle | *Quiscalus major* | 1 | 1 |
| *Icteridae* | Common grackle | *Quiscalus quiscula* | 95 | 1 |
| *Icteridae* | Blackbird (unidentified) | | 4 | 0 |
| *Mimidae* | Gray catbird | *Dumetella carolinensis* | 9 | 0 |
| *Mimidae* | Northern mockingbird | *Mimus polyglottos* | 1 | 0 |
| *Mimidae* | Curve-billed thrasher | *Toxostoma curvirostre* | 1 | 0 |

| | | | | |
|---|---|---|---|---|
| *Mimidae* | Brown thrasher | *Toxostoma rufum* | 1 | 0 |
| *Oriolodae* | Oriole (unidentified) | | 7 | 0 |
| *Paridae* | Tufted titmouse | *Baeolophus bicolor* | 2 | 0 |
| *Paridae* | Black-capped chickadee | *Poecile atricapillus* | 26 | 0 |
| *Parulidae* | Common yellowthroat | *Geothlypis trichas* | 1 | 0 |
| *Parulidae* | Orange-crowned warbler | *Leiothlypis celata* | 1 | 0 |
| *Parulidae* | Magnolia warbler | *Setophaga magnolia* | 15 | 0 |
| *Parulidae* | Yellow warbler | *Setophaga petechia* | 2 | 0 |
| *Parulidae* | Warbler (unidentified) | | 13 | 0 |
| *Passerellidae* | Dark-eyed Junco | *Junco hyemalis* | 19 | 1 |
| *Passerellidae* | Song sparrow | *Melospiza melodia* | 1 | 0 |
| *Passerellidae* | Fox sparrow | *Passerella iliaca* | 2 | 0 |
| *Passerellidae* | Spotted towhee | *Pipilo maculatus* | 1 | 0 |
| *Passerellidae* | White-throated sparrow | *Zonotrichia albicollis* | 7 | 0 |
| *Passeridae* | House sparrow | *Passer domesticus* | 165 | 1 |
| *Picidae* | Northern flicker | *Colaptes auratus* | 11 | 0 |
| *Picidae* | Red-breasted sapsucker | *Sphyrapicus ruber* | 1 | 1 |
| *Picidae* | Yellow-bellied sapsucker | *Sphyrapicus varius* | 1 | 0 |
| *Picidae* | Woodpecker (unidentified) | | 2 | 0 |
| *Sittidae* | White-breasted nuthatch | *Sitta carolinensis* | 1 | 0 |
| *Sittidae* | Nuthatch (unidentified) | | 3 | 0 |
| *Sturnidae* | European starling | *Sturnus vulgaris* | 72 | 0 |
| *Thraupidae* | Tanager (unidentified) | | 3 | 0 |
| *Troglodytidae* | Carolina wren | *Thryothorus ludovicianus* | 11 | 0 |
| *Troglodytidae* | Pacific wren | *Troglodytes pacificus* | 1 | 0 |
| *Troglodytidae* | Wren (unidentified) | | 2 | 0 |
| *Turdidae* | American robbin | *Turdus migratorius* | 170 | 1 |
| *Turdidae* | Eastern bluebird | *Sialia sialis* | 31 | 0 |
| *Turdidae* | Veery | *Catharus fuscescens* | 1 | 0 |
| *Turdidae* | Hermit thrush | *Catharus guttatus* | 2 | 0 |
| *Turdidae* | Swainsons thrush | *Catharus ustulatus* | 2 | 0 |
| *Turdidae* | Wood thrush | *Hylocichla mustelina* | 14 | 0 |
| *Turdidae* | Varied thrush | *Ixoreus naevius* | 20 | 0 |
| *Turdidae* | Townsend's solitaire | *Myadestes townsendi* | 1 | 0 |
| *Turdidae* | Thrush (unidentified) | | 9 | 0 |
| *Tyrannidae* | Eastern phoebe | *Sayornis phoebe* | 12 | 0 |
| *Vireonidae* | Red-eyed vireo | *Vireo olivaceus* | 2 | 0 |
| *Corvidae* | American crow | *Corvus brachyrhynchos* | 301 | 30 |
| *Corvidae* | Common raven | *Corvus corax* | 106 | 25 |
| *Corvidae* | Fish crow | *Corvus ossifragus* | 24 | 5 |

| | | | | |
|---|---|---|---|---|
| *Corvidae* | Blue jay | *Cyanocitta cristata* | 39 | 0 |
| *Corvidae* | Steller's jay | *Cyanocitta stelleri* | 5 | 0 |
| *Corvidae* | Green jay | *Cyanocorax luxuosus* | 3 | 0 |
| *Corvidae* | Gray jay | *Perisoreus canadensis* | 1 | 0 |
| *Corvidae* | Black-billed magpie | *Pica hudsonia* | 35 | 6 |
| *Corvidae* | Crow (unidentified) | | 8 | 0 |
| *Corvidae* | Magpie (unidentified) | | 7 | 0 |
| *Corvidae* | Scrub jay (unidentified) | | 3 | 0 |
| *Odontophoridae* | Northern bobwhite quail | *Colinus virginianus* | 56 | 0 |
| *Odontophoridae* | Quail (unidentified) | | 701 | 3 |
| *Phasianidae* | Ruffed grouse | *Bonasa umbellus* | 15 | 1 |
| *Phasianidae* | Spruce grouse | *Canachites canadensis* | 1 | 0 |
| *Phasianidae* | Greater sage-grouse | *Centrocercus urophasianus* | 5 | 1 |
| *Phasianidae* | Willow ptarmigan | *Lagopus lagopus* | 1 | 0 |
| *Phasianidae* | Wild turkey | *Meleagris gallopavo* | 451 | 29 |
| *Phasianidae* | Ring-necked pheasant | *Phasianus colchicus* | 233 | 31 |
| *Phasianidae* | Sharp-tailed grouse | *Tympanuchus phasianellus* | 40 | 0 |
| *Phasianidae* | Grouse (unidentified) | | 10 | 0 |
| *Phasianidae* | Pheasant (unidentified) | | 674 | 61 |
| *Accipitridae* | Cooper's hawk | *Accipiter cooperii* | 279 | 22 |
| *Accipitridae* | Northern goshawk | *Accipiter gentilis* | 2 | 0 |
| *Accipitridae* | Sharp-shinned hawk | *Accipiter striatus* | 55 | 4 |
| *Accipitridae* | Golden eagle | *Aquila chrysaetos* | 69 | 3 |
| *Accipitridae* | Red-tailed hawk | *Buteo jamaicensis* | 747 | 193 |
| *Accipitridae* | Rough-legged hawk | *Buteo lagopus* | 16 | 8 |
| *Accipitridae* | Red-shouldered hawk | *Buteo lineatus* | 179 | 20 |
| *Accipitridae* | Broad-winged hawk | *Buteo platypterus* | 120 | 6 |
| *Accipitridae* | Ferruginous hawk | *Buteo regalis* | 1 | 0 |
| *Accipitridae* | Swainson's hawk | *Buteo swainsoni* | 19 | 4 |
| *Accipitridae* | Norther harrier | *Circus hudsonius* | 2 | 0 |
| *Accipitridae* | Bald eagle | *Haliaeetus leucocephalus* | 1150 | 294 |
| *Accipitridae* | Harris's hawk | *Parabuteo unicinctus* | 27 | 0 |
| *Accipitridae* | Mississippi kite | *Ictinia mississippiensis* | 11 | 0 |
| *Accipitridae* | Eagle (unidentified) | | 13 | 2 |
| *Accipitridae* | Hawk (unidentified) | | 78 | 10 |
| *Cathartidae* | Turkey vulture | *Cathartes aura* | 186 | 73 |
| *Cathartidae* | Black vulture | *Coragyps atratus* | 495 | 336 |
| *Cathartidae* | California condor | *Gymnogyps californianus* | 80 | 3 |
| *Cathartidae* | Vulture (unidentified) | | 34 | 12 |
| *Falconidae* | Merlin | *Falco columbarius* | 46 | 4 |

| | | | | |
|---|---|---|---|---|
| *Falconidae* | Aplomado falcon | *Falco femoralis* | 2 | 0 |
| *Falconidae* | Prairie falcon | *Falco mexicanus* | 1 | 1 |
| *Falconidae* | Peregrine falcon | *Falco peregrinus* | 148 | 46 |
| *Falconidae* | Gyrfalcon | *Falco rusticolus* | 80 | 0 |
| *Falconidae* | American kestrel | *Falco sparverius* | 101 | 3 |
| *Falconidae* | Falcon (unidentified) | | 9 | 5 |
| *Pandionidae* | Osprey | *Pandion haliaetus* | 82 | 4 |
| *Strigidae* | Northern saw-whet owl | *Aegolius acadicus* | 5 | 0 |
| *Strigidae* | Boreal owl | *Aegolius funereus* | 1 | 0 |
| *Strigidae* | Short-eared owl | *Asio flammeus* | 3 | 1 |
| *Strigidae* | Long-eared owl | *Asio otus* | 11 | 1 |
| *Strigidae* | Burrowing owl | *Athene cunicularia* | 7 | 0 |
| *Strigidae* | Snowy owl | *Bubo scandiacus* | 28 | 9 |
| *Strigidae* | Great horned owl | *Bubo virginianus* | 610 | 234 |
| *Strigidae* | Western screech owl | *Megascops kennicottii* | 5 | 0 |
| *Strigidae* | Eastern screech owl | *Magascops asio* | 96 | 3 |
| *Strigidae* | Flammulated owl | *Psiloscops flammeolus* | 5 | 0 |
| *Strigidae* | Great grey owl | *Strix nebulosa* | 4 | 0 |
| *Strigidae* | Spotted owl | *Strix occidentalis* | 1 | 0 |
| *Strigidae* | Barred owl | *Strix varia* | 320 | 23 |
| *Strigidae* | Northern hawk-owl | *Surnia ulula* | 1 | 0 |
| *Strigidae* | Owl (unidentified) | | 108 | 10 |
| *Strigidae* | Screech owl (unidentified) | | 8 | 0 |
| *Tytonidae* | Barn owl | *Tyto alba* | 61 | 0 |
